# Inferring local molecular dynamics from the global actin network structure: a case study of 2D synthetic branching actin networks

**DOI:** 10.1101/2022.09.06.506753

**Authors:** Minghao W. Rostami, Brittany E. Bannish, Kelsey Gasior, Rebecca L. Pinals, Calina Copos, Adriana T. Dawes

**Affiliations:** Syracuse University; University of Central Oklahoma; University of Ottawa; Massachusetts Institute of Technology; Northeastern University; The Ohio State University

## Abstract

Cells rely on their cytoskeleton for key processes including division and directed motility. Actin filaments are a primary constituent of the cytoskeleton. Although actin filaments can create a variety of network architectures linked to distinct cell functions, the microscale molecular interactions that give rise to these macroscale structures are not well understood. In this work, we investigate the microscale mechanisms that produce different branched actin network structures using an iterative classification approach. First, we employ a simple yet comprehensive agent-based model that produces synthetic actin networks with precise control over the microscale dynamics. Then we apply machine learning techniques to classify actin networks based on measurable network density and geometry, identifying key mechanistic processes that lead to particular branched actin network architectures. Extensive computational experiments reveal that the most accurate method uses a combination of supervised learning based on network density and unsupervised learning based on network symmetry. This framework can potentially serve as a powerful tool to discover the molecular interactions that produce the wide variety of actin network configurations associated with normal development as well as pathological conditions such as cancer.

## 1. Introduction

Actin, the most abundant protein in eukaryotic cells, is involved in a wide range of key cellular functions including cell motility, cell differentiation, muscle contraction, and cytokinesis (1; 2). Actin monomers form rod-like actin polymers, which are constantly assembled, disassembled, and remodeled by accessory proteins and molecular motors. Because of the noise-dominated complex dynamics, the resulting actin networks have a variety of architectures with distinguishable biochemical and mechanical properties. One notable actin meshwork is the branching protrusive network in the lamellipodium, which is the thin, sheet-like extension used for directed cell movement on flat surfaces (3). Previous quantitative studies have demonstrated that the growth of the branching network must be primarily two-dimensional to maintain the integrity of the lamellipodium (4). Abnormalities in the structure and function of the lamellipodium have been implicated in a variety of diseases, including cancer metastasis, neurological disorders, and immune disorders (5). Despite the importance of the lamellipodium in cellular functions, a direct and comprehensive link from local molecular dynamics to network organization to cellular behavior has yet to be established.

Branched actin networks emerge as a result of the complex interplay between various known molecular processes as follows (2). Actin filaments grow or shrink through the addition or loss of actin monomers from their pointed and barbed ends. Filament branching occurs when an Arp2/3 protein complex binds to a filament and creates a nucleation site for a new filament to extend from the existing one at a 70° angle (6). Capping proteins prevent the additional growth of an actin filament by binding to the barbed end. Depending on the intracellular environment that actin networks grow in, including the availability and distribution of resources (actin monomers) and regulators (capping proteins and Arp2/3 complexes), branched actin structures can exhibit vastly different shapes and internal organizations. Our focus is on the inverse problem: given such a network, what are the underlying growth conditions that gave rise to the observed architecture? The power of such a technique lies in its predictive nature; for example, we can predict that a “spiky” network is due to limited availability of Arp2/3 complexes.

To address the need for an inverse mapping from macroscopic networks to underlying microscopic formation dynamics, we present a novel analysis pipeline. The pipeline is a two-pronged approach based on synthetic networks with prescribed local dynamics, along with machine learning and topological tools. The first prong is an agent-based stochastic model that allows us to construct protrusive actin structures in two dimensions. Building on our previously published work, we create a stochastic model that generates actin structures by tuning various interactions and parameters (7). With this *in silico* approach, networks are generated quickly and inexpensively while molecular processes and parameters are controlled in a systematic way. The generation of synthetic data is an important step for our classification method since few experiments exist for lamellipodium-like network growth in controlled environments.

The synthetic branched actin networks are used in the second part of our approach, which applies machine learning techniques to identify the governing principles that give rise to a network architecture. We formulate this process as a classification problem in which we classify images of networks with respect to the underlying local molecular interactions. To solve this problem, we then leverage the data-processing power of a variety of machine learning algorithms including the convolutional neural networks (CNNs), which specialize in processing data on a grid and have been successful at image classification. We perform both fine-grained classification and coarse-grained classification; the former aims to pinpoint the exact combination of molecular processes that give rise to a specific actin structure, whereas the latter uncovers the distinct dominating mechanisms. The coarse-grained classification is developed based on the observation that different mechanisms can lead to very similar network architectures, making fine-grained classification unreliable. Our improved approach combines supervised (CNN) and unsupervised (k-means clustering) learning methods, and utilizes both raw and symmetry-transformed data. Using simulated data, we find that the fine-grained classifier has an accuracy ranging between 66% and 90% depending on the network growth condition, while the coarse-grained classifier boasts an accuracy of 87% − 99%. The unsupervised regrouping of the growth conditions into coarse-grained classes identifies limiting capping proteins and branching complexes as primary drivers of network architecture. The success of our method on synthetic networks is a promising step towards the ultimate discovery and analysis of regulatory mechanisms in experimentally obtained actin networks.

## 2. Methods

### 2.1. Generation of synthetic branching actin networks

In our previous work, we built an agent-based computational framework to capture the local microstructure of branching actin networks under various intracellular conditions (7) (top, Figure 1). The framework uses a molecular-level representation of dynamically assembling and disassembling polymers (actin filaments) in two dimensions. In order to capture more of the biological complexity of branching networks, we extend this framework to include capping and limited molecular resources in addition to polymerization, depolymerization, and branching of polar actin filaments.

**Figure 1:**
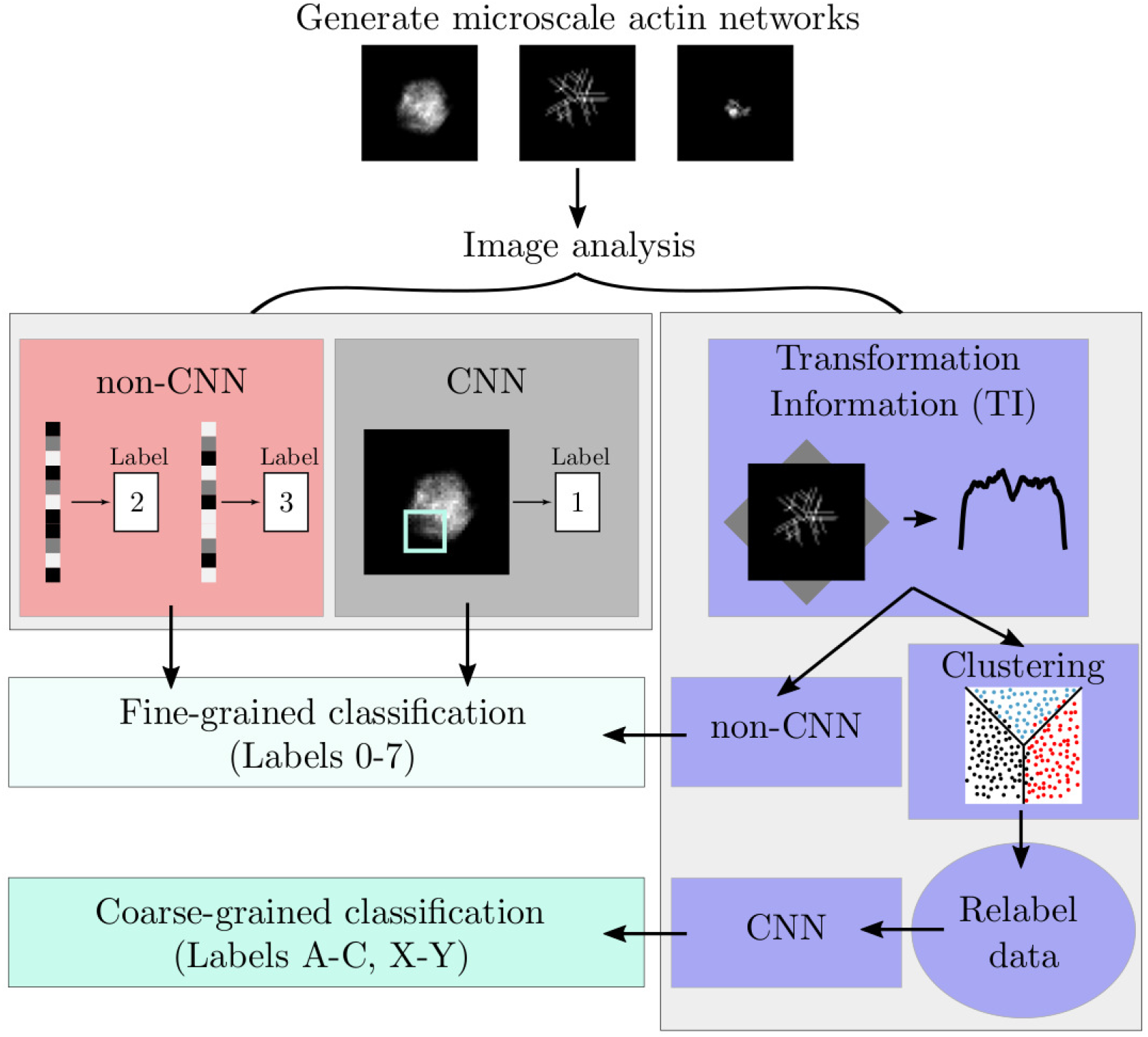
Outline of the analysis pipeline for identifying the governing dynamics of observed architectures of synthetic branched actin networks. A detailed description of the analysis, which combines agent-based modeling, machine learning methods, and image processing, is in Section 2.

The dynamics used to generate synthetic actin networks are summarized in Figure 2A. Implementation of network dynamics and rates is chosen to closely match biological measurements. A branching actin network is formed from a single nucleation site in a two-dimensional domain. By excluding the third spatial dimension, we assume the branching actin network to be relatively flat as observed in lamellipodia (8). Filaments are represented as rigid rods with a spatially fixed base and a barbed tip capable of growing or shrinking. Changes in both actin filament length and overall network structure are due to the addition or removal of single actin monomers and branching via branching proteins (Arp2/3 nucleation complexes) at the free end. Filament growth and shrinkage occur in discrete increments corresponding to the length of an actin monomer, 0.0027 *μ*m (9). Based on the known effect of the Arp2/3 complex, the newly branched filament is assigned to grow in a direction that is 70° on either side of the preexisting filament tip (10). Capping proteins are incorporated and they regulate actin filament length by binding to the barbed end of a filament; this blocks the addition or loss of monomers from that filament (11). The concentration of free monomers and branching proteins can be either unlimited or limited; both cases are considered in our work. Additional implementation details are available in Appendix A.1, along with model parameters and computational constants found in Table A1 and explained in Appendix A.2. The simulation and analysis of actin networks are implemented in a custom Matlab code.^2^

**Figure 2:**
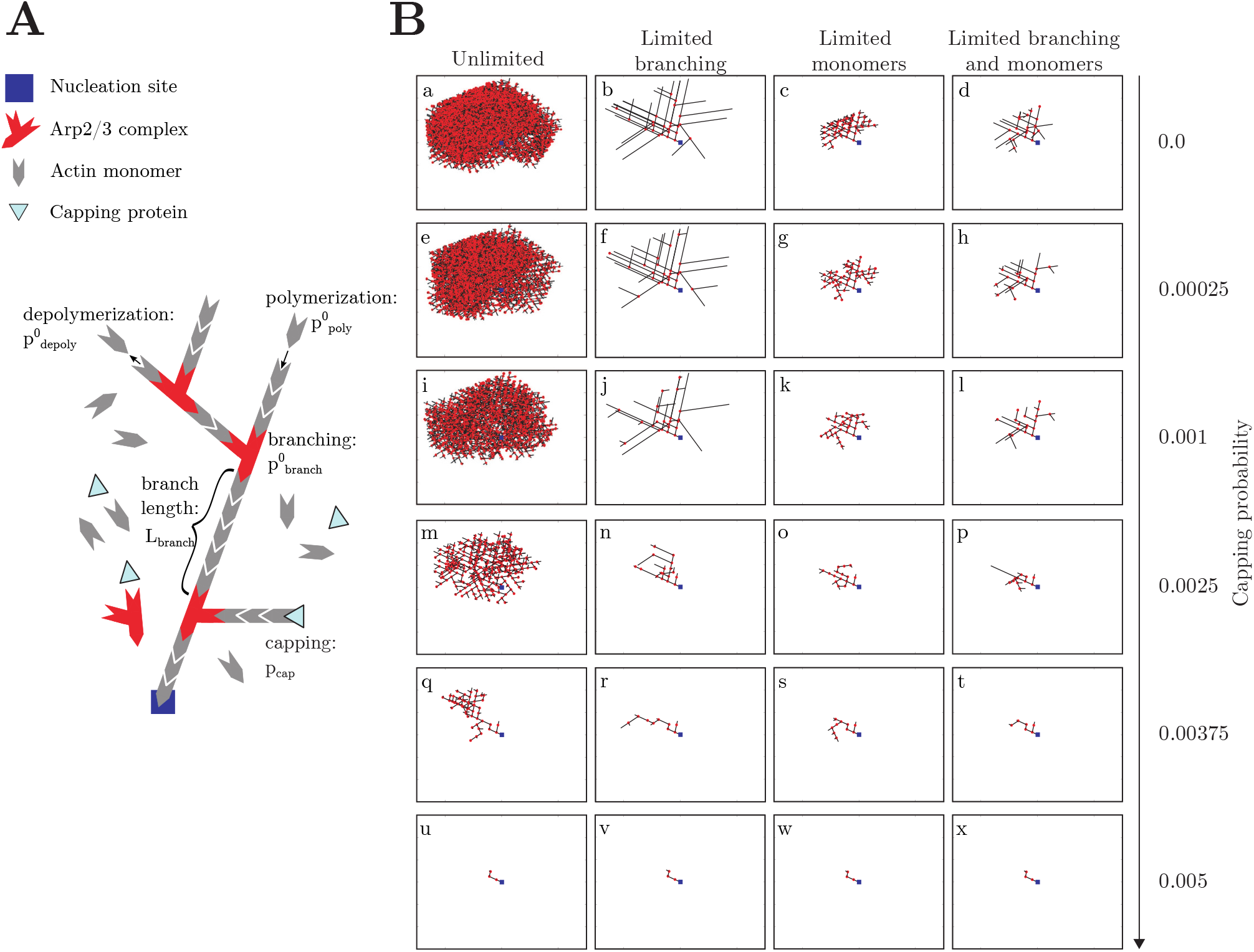
(A)Schematic of actin kinetics incorporated in our simulation framework. Actin filament growth begins at the initial nucleation site (blue) through polymerization. Monomers have a polymerization probability of 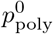 and a depolymerization probability of 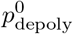. Attachment of Arp2/3 complex (red) to the filament facilitates Y-branching. Association or disassociation of monomers to/from a filament can be halted by the binding of capping proteins (light blue) with probability *p*_cap_. (B) **Sample actin network architectures arising from various growth conditions**. All networks are shown after 10 s with a simulation domain of a 10 *μ*m × 10 *μ*m square. For ease of comparison, the initial angle of growth from the nucleation site was kept constant between all simulations.

The spatial domain of each simulation is a 10 *μ*m × 10 *μ*m square, which is comparable to the size of eukaryotic cells. To extrapolate a corresponding density from a discrete actin network architecture, the computational domain is uniformly subdivided into a 49 × 49 grid of size 0.2 *μ*m × 0.2 *μ*m. The choice of grid resolution represents a balance between computational cost and underlying filament and monomer sizes. At the end of a given simulation, we calculate the total length of actin filaments contained in each discretized grid box. Since this length is a scalar multiple of the number of actin monomers contained in a grid box, we take the length divided by the area of the grid box, 0.04 *μ*m^2^, to be the actin density at the center of that box. The result is a 49 × 49 grayscale image of actin densities, where the brighter a pixel is, the higher the actin density is at that location (see the top panel of Figure 1 and Appendix A.3).

The data used to train and validate the classification algorithms consists of density images of actin networks generated under eight possible growth conditions labeled 0, 1, 2,…, 7. These eight categories include all combinations of capping or no capping, limited or unlimited Arp2/3 branching complexes, and limited or unlimited actin monomers (Table 1). For each of the six capping probabilities considered (Table 2) and each of the eight growth conditions, 300 independent simulations are run, totaling 2400 simulated actin networks to be analyzed and classified for their underlying molecular dynamics. The networks are divided into training, validation, and test sets as follows. A group of 75 networks for each growth condition (600 networks total) are randomly selected for the training data. From the remaining networks, we randomly select a group of 25 networks per condition (200 networks total) to be used for validation. The networks that have not been selected for either the training set or validation set (200 per condition, 1600 networks total) comprise the test data. The training, validation, and test sets are disjoint and produced by independent runs of the microscale model.

**Table 1:** Fine-grained labels. Eight possible fine-grained growth conditions for actin networks.

| Label | Capping | Branching | Monomers |
| --- | --- | --- | --- |
| 0 ● | × | Unlimited | Unlimited |
| 1 ● | ✓ | Unlimited | Unlimited |
| 2 ● | × | Limited | Unlimited |
| 3 ● | × | Unlimited | Limited |
| 4 ● | ✓ | Limited | Unlimited |
| 5 ● | ✓ | Unlimited | Limited |
| 6 ● | × | Limited | Limited |
| 7 ● | ✓ | Limited | Limited |

**Table 2:** Accuracy of all tested machine learning classification algorithms of synthetic actin network architectures.

| Capping probability | non-CNN (best) | Fine-grained CNN | TI-based non-CNN (best) | Coarse-grained CNN |
| --- | --- | --- | --- | --- |
| 0.00025 | 69% (quadratic SVM) | 82% | 53% (trilayered NN) | 99% |
| 0.00050 | 84% (linear SVM) | 90% | 64% (linear SVM) | 98% |
| 0.00100 | 81% (quadratic SVM) | 89% | 66% (quadratic SVM) | 96% |
| 0.00150 | 77% (medium Gaussian SVM) | 84% | 63% (median NN) | 87% |
| 0.00250 | 67% (narrow NN) | 74% | 61% (narrow NN) | 87% |
| 0.00375 | 65% (medium Gaussian SVM) | 66% | 52% (quadratic SVM) | 98% |

### 2.2. “Fine-grained” actin network classification

To assign a class or label to a given actin network as resulting from one of the eight growth conditions, we build classifiers using supervised machine learning techniques based on the images of actin densities and compare their performance. Foreshadowing the findings in the next section, we refer to this type of classification as “fine-grained.”

#### 2.2.1. Classification based on density

The supervised machine learning algorithms considered are: convolutional neural network (CNN) (12), support-vector machine (SVM) (13), *k*-nearest neighbors (k-NN) (14), ensemble algorithms (15; 16), and neural network (NN) (17). We use “non-CNN” to mean supervised machine learning methods other than CNN. All classifiers are built for the two-dimensional actin density plots taken at a fixed time point as described in Section 2.1. The CNN classifier uses the density data in its natural dimension, a 49 × 49 matrix (image), whereas the non-CNN classifiers use the same data arranged into a 49^2^ 1 vector. For each of the six capping probabilities considered (see Table 2), we proceed as follows:

**Step 1** Generate training data: The training data is composed of density plots of 600 actin networks (75 networks per growth condition) simulated by the same number of independent runs of the microscale model and labeled by their classes 0, 1, 2, …, 7 according to their respective growth conditions.

**Step 2** Train classifiers: We consider CNN and 22 non-CNN classifiers:

- six SVM variants: linear, quadratic, cubic, fine Gaussian, medium Gaussian, and coarse Gaussian;
- six k-NN variants: fine, medium, coarse, cosine, cubic, and weighted;
- five ensemble algorithms: boosted trees, bagged trees, subspace discriminant, sub-space k-NN, and random undersampling boosted trees;
- five NN variants: narrow, medium, wide, bilayered, and trilayered.

Matlab’s Classification Learner App^3^ is used for all non-CNN classifiers and the CNN classifier is implemented using a custom code with built-in functions from Matlab’s Deep Learning Toolbox ^4^ and Statistics and Machine Learning Toolbox^5^. All 23 classifiers are trained on networks in the training set. Once trained, each classifier is a function that maps the density image (for the CNN classifier) or vector (for a non-CNN classifier) of a network to its predicted label, an integer between 0 and 7.

**Step 3** Evaluate performance: We calculate and compare the accuracy of all classifiers on data not used in the training process. A classifier is successful at labeling an actin density plot if its predicted label agrees with the true label and unsuccessful otherwise. The percentage accuracy of a classifier is defined as

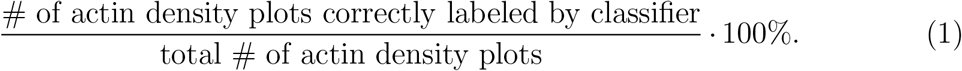

First, we calculate the percentage accuracy of the 22 non-CNN classifiers on the validation data composed of 200 images of actin density (25 per growth condition). This allows us to identify the best non-CNN classifier as the one with the highest percentage accuracy. Then, to compare the best performing non-CNN classifier and the CNN classifier, we compute their respective percentage accuracy on the test data, which consists of 1600 actin density plots (200 per condition).

#### 2.2.2. Quantification of shape symmetry

To identify geometric properties of simulated actin networks, we calculate the transformation information (TI) associated with each network density plot as outlined in (18). The TI function quantifies the amount of information lost by approximating the original image by the transformed image. More specifically, the value of the TI function measures how approximate the symmetry is, with minima of the TI function indicating the transformation that results in the least change from the original image. Here, an affine transformation corresponding to rotation is used to quantify the symmetry properties in the actin network. The function *μ* (**x**)is the actin density at location **x** in each image. The steps for calculating the TI curve are outlined below and shown graphically in Figure 3:

**Figure 3:**
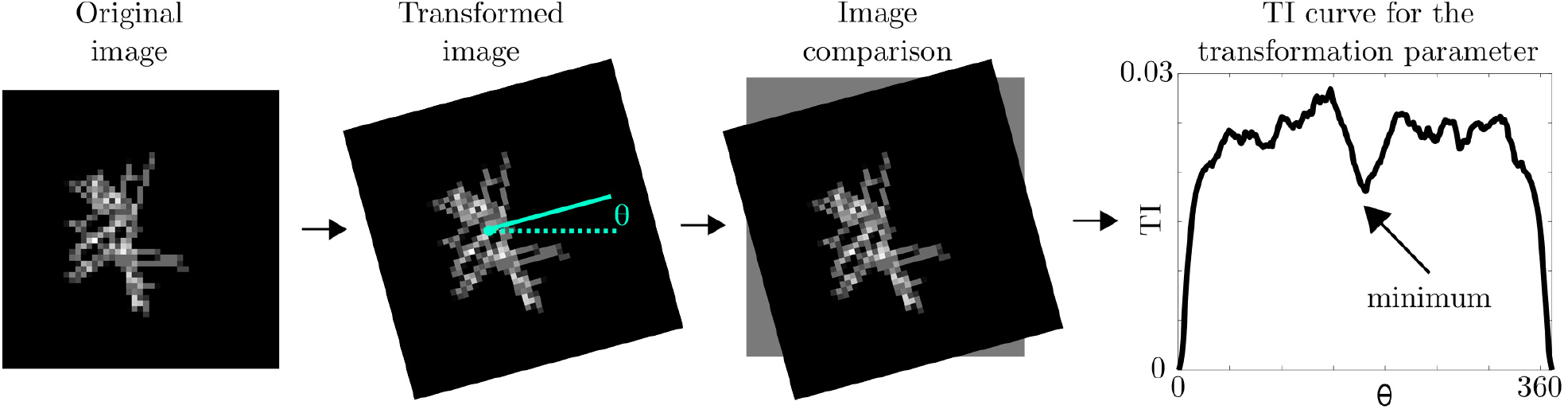
Schematic for quantification of approximate symmetries in the actin density images using transformation information (TI). The image is modified using an affine transformation (in this case, rotation by angle *θ*). The original and transformed images are compared to determine the amount of information lost by using the newly transformed image to approximate the original. Minima of the TI curve indicate approximate symmetries in the actin network.

**Step 1** Obtain TI measurements by:

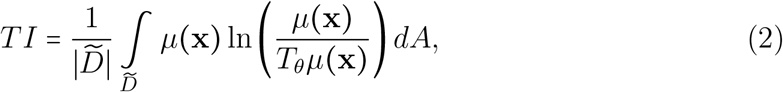

where *T*_*θ*_ is the rotation of the image by angle *θ*, 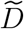 is the domain intersection of the original and transformed images, 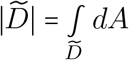, and *dA* is the area element. We repeat the calculation and compute the TI for each increment of the transformation parameter, angle *θ*, to obtain TI as a function of angle *θ*.

**Step 2** Identify approximate symmetries: Find and rank minima of the TI curve in order to identify approximate symmetries of the actin density plots. For example, a TI value of 0 indicates perfect symmetry.

Further details of the TI calculation and Matlab code are in (18).

#### 2.2.3. Classification based on shape symmetry

For each plot of the actin network density, the TI calculation produces a 361 × 1 vector (or a curve); the *i*^th^ entry of the vector is the TI measurement for the transformation parameter *θ*_*i*_ = (*i* − 1) degrees for *i* = 1, 2, …, 361. The non-CNN supervised classifiers can also be applied to the TI curves extracted from the actin density plots. The process is the same to the one described in Section 2.2.1 since the TI curves carry the same fine-grained labels 0, 1, …, 7 associated with growth conditions.

### 2.3. “Coarse-grained” actin network classification

Different growth conditions may result in similar network architectures. Therefore, fewer, more identifiable classes are sometimes desired. In order to reveal governing motifs of emergent network architectures, we propose a “coarse-grained” classification which results in fewer than eight classes. Our approach combines supervised and unsupervised machine learning techniques; first, an unsupervised algorithm is applied to identify the new, fewer than eight, groupings in the dimension-reduced TI curves extracted from actin density plots, and then a supervised learning algorithm is used to build a classifier that determines which new grouping (or label) to assign to a particular actin density plot. The procedure is outlined below.

**Step 1** Identify new classes: To abstract away fine-level detail, we use principle component analysis (PCA) to transform each TI curve (a 361 × 1 vector) to a 2 × 1 vector referred to as the “reduced” TI curve. The reduced TI curve consists of the two leading coefficients given by PCA. Next, we apply k-means, a widely used unsupervised clustering algorithm, to identify the *k* groupings (clusters) of the reduced TI curves (19). We use Matlab’s built-in function for k-means^6^ and repeat it for *k* between 1 and 12. The sum of distances between data points and their group centroids is used to assess the “tightness” of a grouping. Among the twelve values of *k* considered, we choose a relatively small *k* < 8 that is still large enough for the *k* clusters to be identifiable. We then proceed to divide the original eight growth conditions into *k* groups. Growth condition *i* is placed in the *j*th group if the majority of the reduced TI curves originally labeled with *i* are placed in the *j*th group by k-means.

**Step 2** We relabel each actin density plot according to the correspondence between the original eight growth conditions and the new *k* groups identified in Step 1. As an example, consider the case where *k* = 3 and the three groups are labeled A, B, C. If the original label 4 is deemed a member of the group A in Step 1, then all the density plots previously labeled with 4 will be relabeled with A.

**Step 3** Rebuild and re-evaluate the CNN classifier: Repeat Steps 2 to 3 in Section 2.2.1 using the relabeled data.

## 3. Results

We present a robust, rotation-invariant, and noise-insensitive computational pipeline for the inverse mapping from a synthetic actin network to its underlying molecular kinetics. In the pipeline, an actin network is fed through the fine-grained classifier to identify molecular growth conditions (Sections 2.2.1 and 2.2.3) or through the coarse-grained classifier to identify primary dynamics (Section 2.3). Training the supervised statistical methods for accurate classification may require a large set of networks labeled according to their underlying molecular kinetics. To rapidly generate tens of thousands of actin networks with known dynamics, we employ our molecular kinetics-based modeling framework outlined in Section 2.1.

### 3.1. Stochastic simulations produce synthetic actin networks with a variety of architectures, particularly in the presence of limited resources

We adjust parameter values in the stochastic simulations to generate diverse actin networks, some of which are shown in Figure 2B. For the networks presented here, the polymerization and depolymerization probabilities are held constant across simulations, while the capping probability is varied (see Appendix A.2 for details).

As expected, the addition of capping proteins and limited resources to the actin kinetics modeling framework, individually or in combination, changes the resulting network architecture (Figure 2B). With unlimited resources and no capping, a dense and highly branched network forms (Figure 2B.a). The network remains dense and branched for low capping probabilities (*p*_cap_ = 0.00025 and *p*_cap_ = 0.001), but as the capping probability increases further,polymerization and depolymerization are halted, resulting in much smaller, sparser networks (Figure 2B.m,q,u). In cases with low or no capping, the addition of limited Arp2/3 branching complexes (Figure 2B.b,f,j) results in longer filaments once all the Arp2/3 complexes are used; with no additional Arp2/3 complexes, no further branching can occur, so filaments only polymerize and depolymerize. These longer filaments are not seen at higher capping probabilities (Figure 2B.n,r,v) because filaments are capped before they can grow too long. Regardless of the capping probability, simulations with limited monomers result in significantly smaller networks (Figure 2B.c,g,k,o,s,w); this is because once all the actin monomers are used, no further bulk growth can occur and only slight structural modifications to the network continue, as individual monomers depolymerize and re-polymerize elsewhere in the network. Finally, in cases with both limited Arp2/3 complexes and limited monomers (Figure 2B.d,h,l,p,t,x), networks are spatially reduced, with emergence of some longer filaments at low capping probabilities (Figure 2B.d,h,l,p) that are not present at higher capping probabilities (Figure 2B.t,x).

### 3.2. Extreme capping rates obscure the role of other dynamics when classifying network structures using standard machine learning techniques

Our first attempt at constructing a classifier relies on non-CNN supervised machine learning algorithms: support-vector machines, k-nearest neighbors, ensemble algorithms, and neural networks. For each capping probability listed in Table 2, we generate 300 actin density plots for each of the eight growth conditions with labels *j* = 0, 1, 2, …, 7. The generated data is partitioned into training, validation, and test sets as described in Section 2.1. The leftmost three columns in Figure 4 show 12 actin density plots, with their respective labels placed above them, randomly selected from the training set for an intermediate capping probability, 0.001. The accuracy of each algorithm is evaluated on the validation set to determine which classifier has the highest accuracy. The accuracy of this classifier for the test data is reported in Table 2 for each of the six capping probabilities considered. We find that the accuracy of the best non-CNN classifier ranges between 65% and 84% depending on the capping probability. Interestingly, the highest accuracy is obtained with mid-range capping probabilities (0.0005 − 0.0015). That is, at extreme capping rates, the underlying dynamics are not fully uncovered by any of the non-CNN classifiers.

**Figure 4:**
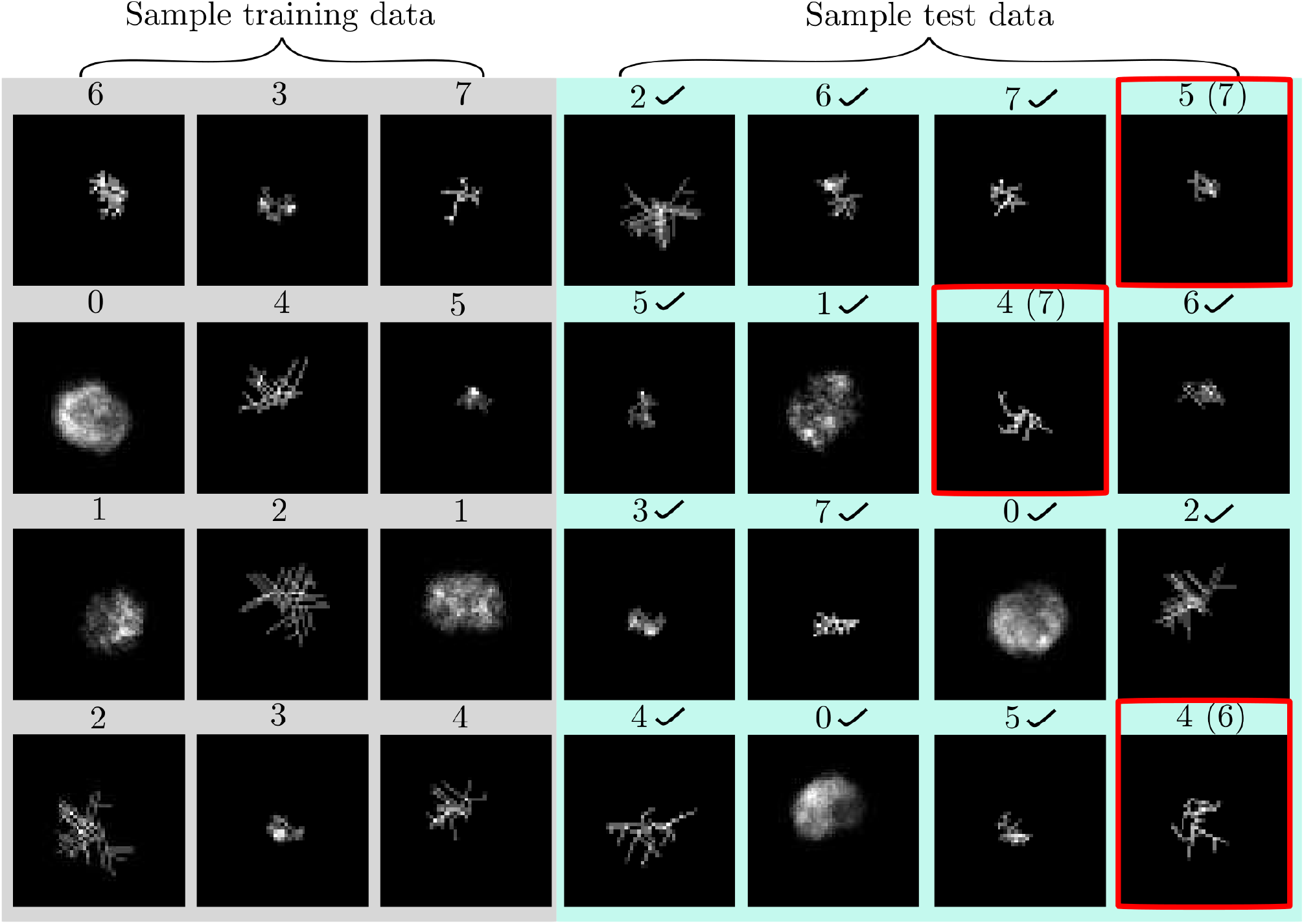
Randomly selected actin density plots and their corresponding fine-grained labels. A brighter pixel indicates a higher actin density and brightness is scaled across an individual figure. Across all images, the capping probability is held fixed at an intermediate value (*p*_cap_ = 0.001). The 12 plots (left, gray) are part of the training set used for tuning the classifiers. The 16 images (right, blue) are from the test set used for evaluating and comparing the classifiers. A check mark indicates that the CNN classifier accurately predicts the true label. The density plots wrongly classified by CNN are outlined in red. For any incorrect CNN predictions, the true labels are listed in parentheses.

### 3.3. Networks grown under widely varying conditions are distinguishable when both actin density and network geometry are evaluated

To improve the overall accuracy of the classification pipeline in the previous section, we turn our attention to CNNs. This class of statistical learning methods has been highly successful at detection of visual imagery (12; 20). We train and evaluate the performance of a CNN classifier as implemented in Section 2.2.1. The validation data is used to inform the choice of CNN hyper-parameters such as the number of network layers and network nodes. The accuracy of the CNN classifier on the test set is summarized in Table 2. The accuracy ranges between 66% and 90%, with an improvement over the best non-CNN classifier for every capping probability. The CNN classifier is more accurate when the capping probability is not too high. In the rightmost four columns in Figure 4, for a fixed capping probability of 0.001, we show 16 actin density plots randomly selected from the test set, along with their true labels and the labels predicted by the CNN classifier if the two are different. For visualization purposes only, the raw data is re-scaled such that for an individual image, the highest density is 1 and the lowest density is 0 (see Appendix A.3 for more details).

Additionally, we find that the CNN classifier is insensitive to rotations (see Appendix A.4) or additive noises up to 10% (see Appendix A.5). Due to the dynamical growth of the network from a nucleation site, the observation time does impact the accuracy of the classification (see Appendix A.6). We observe that if the time at which the test data are collected deviates from the time at which training data are collected by no more than 5%, then the accuracy of the classifier will hardly degrade, allowing room for error in a lab setting where precise control of data collection time is difficult to achieve. As the two times move further apart, however, the accuracy of the classifier can decrease considerably.

Since the CNN classifier uses image data (rather than vectorized data) and hence preserves spatial information of the simulated actin networks, the better performance of the CNN classifier over the non-CNN classifiers suggests that the geometric properties of the network may be important for classification. For higher capping probabilities such as 0.00375, none of the classifiers, CNN or non-CNN, can uncover the growth condition with high accuracy. We speculate that this is because when the capping probability is high, actin networks generated under different growth conditions can approach a similar morphology.

### 3.4. Symmetric properties of actin networks alone are insufficient to distinguish underlying generating mechanisms

To test whether geometric properties of the actin networks alone may accurately classify networks, we use a novel and robust measure of symmetry that is grounded in concepts of entropy and information theory called transformation information (TI) (18). We apply this method to quantify asymmetries of actin density plots under image transformations parameterized by the rotation angle *θ* as outlined in Section 2.2.3 (Figure 3). The outcome is a TI curve associated with each network as a function of angular rotation of the network.

Next, we apply the same standard non-CNN machine learning techniques from Section 3.2 to the TI curves extracted from the same data. That is, we train and evaluate the performance of 22 non-CNN classifiers on TI curves of actin networks grown under the six capping probabilities considered previously. For each capping probability, we identify the best non-CNN classifier as in Section 3.2 and report its accuracy in Table 2. The accuracy ranges between 52% and 66%, lower than density-based classification. Like the density-based classifiers, it performs worse for the extremes of the capping probability range. The best symmetry-based non-CNN classifier is always less accurate than the best density-based non-CNN classifier, indicating that symmetry of the network alone is insufficient to identify the distinct growth condition of an actin network.

To investigate the potential role of network symmetry, we would like to visualize the TI curves but cannot easily do so due to the large dimension, 361, of the data (see Section 2.2.3). A dimension reduction by linear principal component analysis (PCA) is performed on the TI curves. The first two principal components of the linear PCA of the TI curves are calculated, and they account for at least 93% of the variance in the TI data. The visualization of the symmetry information in a reduced two-dimensional space in Figure 5 leads us to hypothesize that the symmetry information can be used to identify coarser level categorization of the data. For example, for the case of low capping probability in Figure 5A, networks with labels 2 (blue) and 5 (magenta) appear to have a similar “vascular” topology. Likewise, networks with labels 0 (red) and 1 (green) share a similar “round” topology, while networks with label 6 (black) are relatively small. Similar conclusions can be drawn for cases with intermediate and high capping probabilities in Figure 5B and 5C respectively.

**Figure 5:**
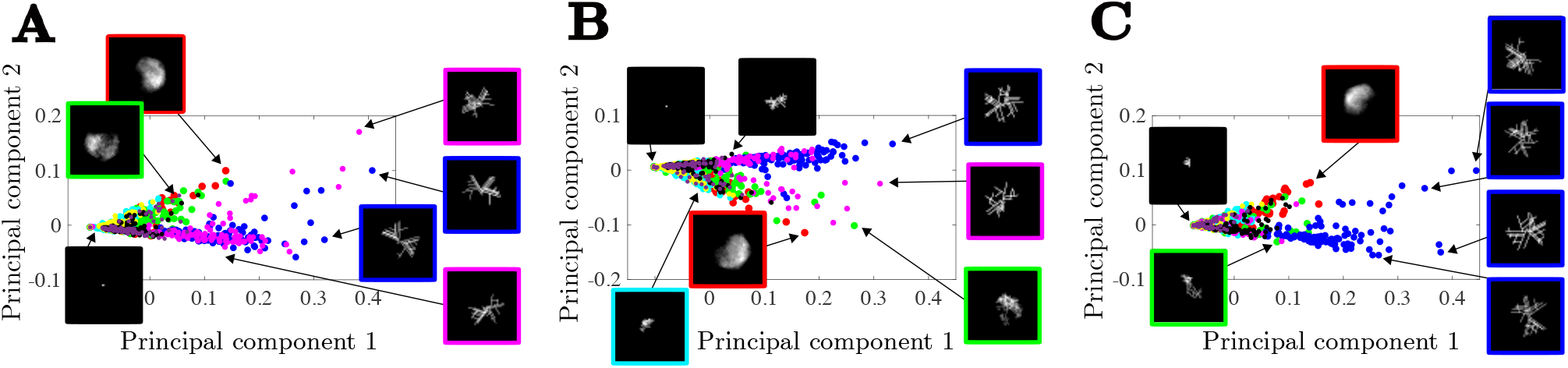
Visualization of the reduced TI curves. Principal components capture key shapes corresponding to “round” versus “spiked-edge” networks. Each dot represents an individual image, and representative images are indicated. Label 0: red; label 1: green; label 2: blue; label 3: cyan; label 4: magenta; label 5: yellow; label 6: black; label 7: purple. (A) Low capping probability, *p*_cap_ = 0.00025. (B) Intermediate capping probability, *p*_cap_ = 0.001. (C) High capping probability, *p*_cap_ = 0.00375.

### 3.5. Symmetry-informed unsupervised clustering reveals three dominant network architecture groupings

Based on the visualization of the dimensionally reduced TI curves of actin density plots, we postulate that there exists a smaller set of dominant dynamics contributing to a given network architecture. We apply the k-means unsupervised clustering algorithm to the reduced TI vectors extracted from the training set to identify a partitioning of the data into *k* groups, where *k* is an integer between 1 and 12. In Figure 6A, for three representative capping probabilities (0.00025, 0.001, and 0.00375), we show the sum of distances between each reduced TI vector and the centroid of the group that it belongs to as the number of groups increases from *k* = 1 to *k* = 12. These plots suggest that *k* = 3 achieves a good balance between the tightness of the groups and the number of groups. The subplots in Figure 6B show the three groups of reduced TI vectors partitioned by k-means for each capping probability. The grouping appears to be predominantly along the first principal component. For capping probabilities of 0.00025 and 0.001, we label the three clusters as “A”, “B”, and “C” and the correspondence between the eight original labels and the three new labels is in Table 3. For low and intermediate capping probabilities, whether and which resources are limited are the main factors in determining the network architecture. Growth conditions with unlimited resources are labeled “A”, growth conditions with limited Arp2/3 complexes and unlimited monomers are labeled “B”, and growth conditions with limited actin monomers are labeled “C”.

**Table 3:** Coarse-grained labels. Three coarse-grained groups for actin networks with low and intermediate capping probability.

| New label | Old label | Capping | Branching | Monomers |
| --- | --- | --- | --- | --- |
| A ▲ | 0 | ✗ | Unlimited | Unlimited |
| A ▲ | 1 | ✓ | Unlimited | Unlimited |
| B ▲ | 2 | ✗ | Limited | Unlimited |
| B ▲ | 4 | ✓ | Limited | Unlimited |
| C ▲ | 3 | ✗ | Unlimited | Limited |
| C ▲ | 5 | ✓ | Unlimited | Limited |
| C ▲ | 6 | ✗ | Limited | Limited |
| C ▲ | 7 | ✓ | Limited | Limited |

**Figure 6:**
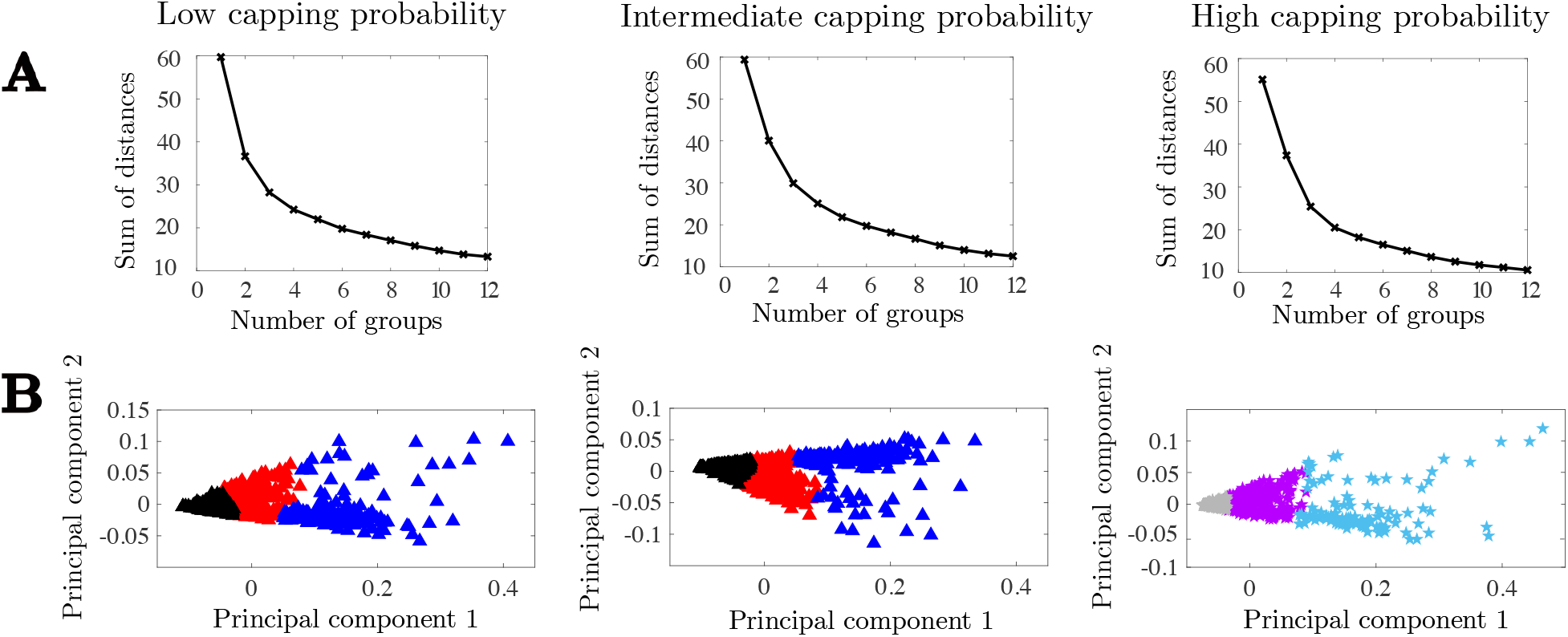
Identification of coarse-grained labels through unsupervised clustering of the reduced TI curves. (A) The sum of the distances between the reduced TI vectors and their respective group centroid, as a function of the number of groups used in the k-means algorithm. (B) Partition of the reduced TI vectors into three groups by k-means. Each triangle or star represents the reduced TI vector extracted from an actin density plot. Label A: red triangle; label B: blue triangle; label C: black triangle; label X: purple star; label Y: gray star; label Z: light blue star. Low, intermediate, and high capping probabilities correspond to *p*_cap_ = 0.00025, 0.001, and 0.00375, respectively.

In the case of high capping probability, we label the three clusters as “X”, “Y”, and “Z” and Table 4 gives the correspondence between the eight fine-grained labels and these three new labels. Whether capping is present and whether branching is limited are the main contributors to the network architecture. Growth conditions that do not include capping are labeled “X” or “Z”, while growth conditions that do include capping are all labeled “Y”. As only one growth condition without capping (the condition which has limited Arp2/3 complexes and unlimited actin monomers) is labeled by “Z”, we can conclude that while capping is the biggest determinant in labeling the data, limited branching is also a feature in distinguishing network architecture.

**Table 4:** Coarse-grained labels at high capping probability. Three coarse-grained groups for actin networks with high capping probability.

| New label | Old label | Capping | Branching | Monomers |
| --- | --- | --- | --- | --- |
| X ★ | 0 | × | Unlimited | Unlimited |
| X ★ | 3 | × | Unlimited | Limited |
| X ★ | 6 | × | Limited | Limited |
| Y ★ | 1 | ✓ | Unlimited | Unlimited |
| Y ★ | 4 | ✓ | Limited | Unlimited |
| Y ★ | 5 | ✓ | Unlimited | Limited |
| Y ★ | 7 | ✓ | Limited | Limited |
| Z ★ | 2 | × | Limited | Unlimited |

### 3.6. Actin networks with defined microscale dynamics can be grouped into three classes with high accuracy even for extreme capping rates

A CNN classifier for coarse-grained classification is constructed, which labels actin networks according to their dominant mechanisms rather than their original growth conditions. The classifier still takes images of actin density and produces predicted labels; the difference is that the labels now correspond to the clusters found by k-means for *k* = 3. The same training and testing data sets are used as previously. In Table 2, the accuracy of the new TI-informed CNN classifier on the test set is reported for the six capping probabilities. We note that the dominant mechanisms can vary with the capping probability, as shown in Section 3.5. For every capping probability considered, the TI-informed CNN classifier is more accurate at the coarse-grained classification than the original CNN classifier is at the fine-grained classification. Now, its accuracy ranges between 87% and 99%, and interestingly, it performs exceedingly well for extreme capping probabilities, a regime where the original CNN classifier has lower accuracy for fine-grained classification. Lastly, in Figure 7, for intermediate capping probability of 0.001, we re-plot the 12 actin networks from the training set and 16 actin networks from the test set shown previously in Figure 4 but now with their coarse-grained labels. The labels predicted by the TI-informed CNN classifier are also included if they are different from the correct coarse-grained labels.

**Figure 7:**
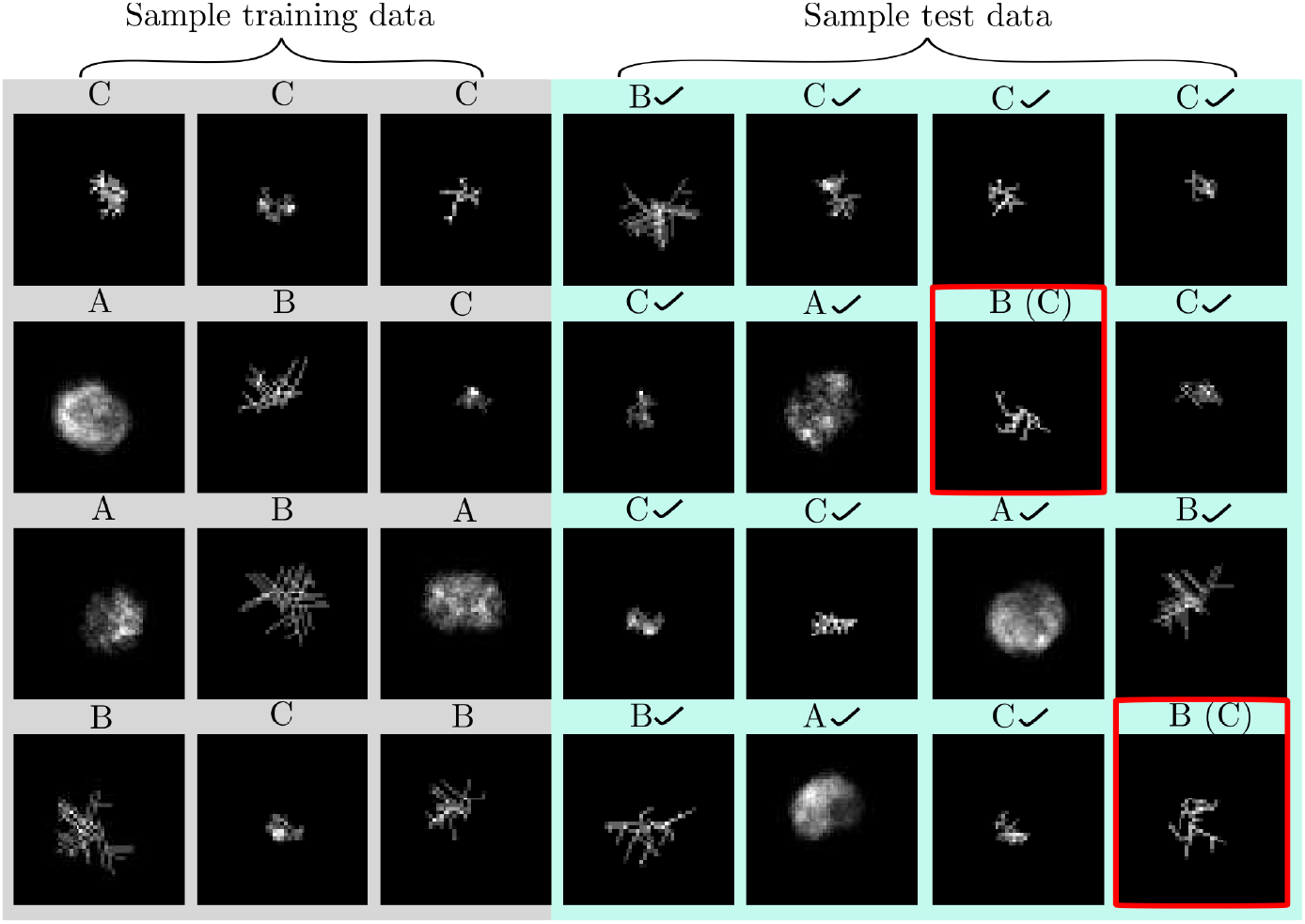
Networks from Figure 4, with their corresponding coarse-grained labels. The images from the test set misclassified by the TI-informed CNN classifier are outlined in red. Their labels predicted by the CNN are listed in parentheses. A check mark indicates that the CNN-predicted label matches the true label. The plots shown are for an intermediate capping probability, *p*_cap_ = 0.001.

## 4. Conclusions & Discussion

Actin is an abundant protein that organizes into network structures that play critical roles in cellular processes, including division and directed cell migration. Defects in actin networks, such as the branching protrusive network in the lamellipodium, are correlated with disease states (21), and studies of actin-related primary immunodeficiencies have also pointed to the mounting evidence of the role of actin structures in a successful immune response (22). Despite the importance of actin network organization, the significance of particular microscale interactions in shaping branched actin networks is not yet clear. Part of the challenge is due to the difficulty of growing branched networks *in vitro* under controlled conditions. Toward this end, we develop an iterative process for network classification: starting with a stochastic model that allows us to “grow” actin networks in a wide variety of settings, we then classify the resulting networks using machine learning techniques informed by network theory-inspired measures. These techniques identify the dominant mechanisms leading to a given actin network, providing mechanistic information and testable hypotheses.

Our stochastic model is able to effectively and efficiently produce a large number of actin networks with known, diverse growth conditions, which are necessary to train our classifiers. The machine learning algorithms in turn identify a small number of meaningful categories hidden in the vast amount of training data, even in the presence of noise and rotations, and allow us to extract information about the dominant microscale dynamics responsible for a given actin network whose growth condition is unknown. By incorporating information from multiple sources (network density and geometry) and focusing on dominant dynamics, we are able to improve classification performance when compared to the same algorithm that uses a single information source (density) and aims to identify detailed dynamics.

We consider a variety of machine learning algorithms based on network density and/or network symmetry and find that the most successful approach is the following: first use an unsupervised clustering algorithm of the approximate network symmetries to identify a few, significant groups of molecular processes, and then use a CNN classifier that has been trained on density data re-labeled according to the groups emerging from unsupervised clustering to classify synthetic networks. CNN has been successfully applied to image analysis, and since actin density is sampled on a two-dimensional grid, CNN is a sensible method to use. Indeed, we observe that CNN is more accurate at network classification than other non-CNN methods considered including the SVMs. This implies that the geometry of actin networks, not just the density, is important in identifying their growth conditions. However, we also find that even with CNN, classifying actin networks according to their precise growth conditions is still challenging since multiple growth conditions can lead to similar actin networks. The importance of network geometry and the desire to accurately identify dominant molecular dynamics motivate our approach of coupling TI-based clustering and CNN classification based on density data.

We use relatively standard algorithms for classification such as CNN, and do not rely on state-of-the-art algorithms for image detection (23), demonstrating that the impact of microscale growth conditions can be distinguished from macroscale images. While it is conceivable that the performance of our methodology could be improved with more sophisticated algorithms, we achieve high accuracy results with CNN. Although we employ synthetic networks, our classification workflow can have broader applications: an experimentalist imaging *in vitro* actin network formation may observe very different network architectures and may wish to identify the biological growth dynamics that led to the given architecture (Figure 8). The tools described here serve as an important step toward that goal. Even in our limited case of a single nucleating site, the discovered labels can provide insight into the growth condition of the network, for example, whether Arp2/3 branching complexes are limited. This prediction can be used in conjunction with perturbations and other data to elucidate the dynamics of the system. In some disease states like cancer, actin network growth is disrupted, and the method described here would be a way to use images of the network architecture to diagnose possible causes of the disrupted growth (24). The analysis framework developed here can be applied more broadly to understand analogous network-forming biological systems, including extensions of fungal hyphae or blood vessels networks.

**Figure 8:**
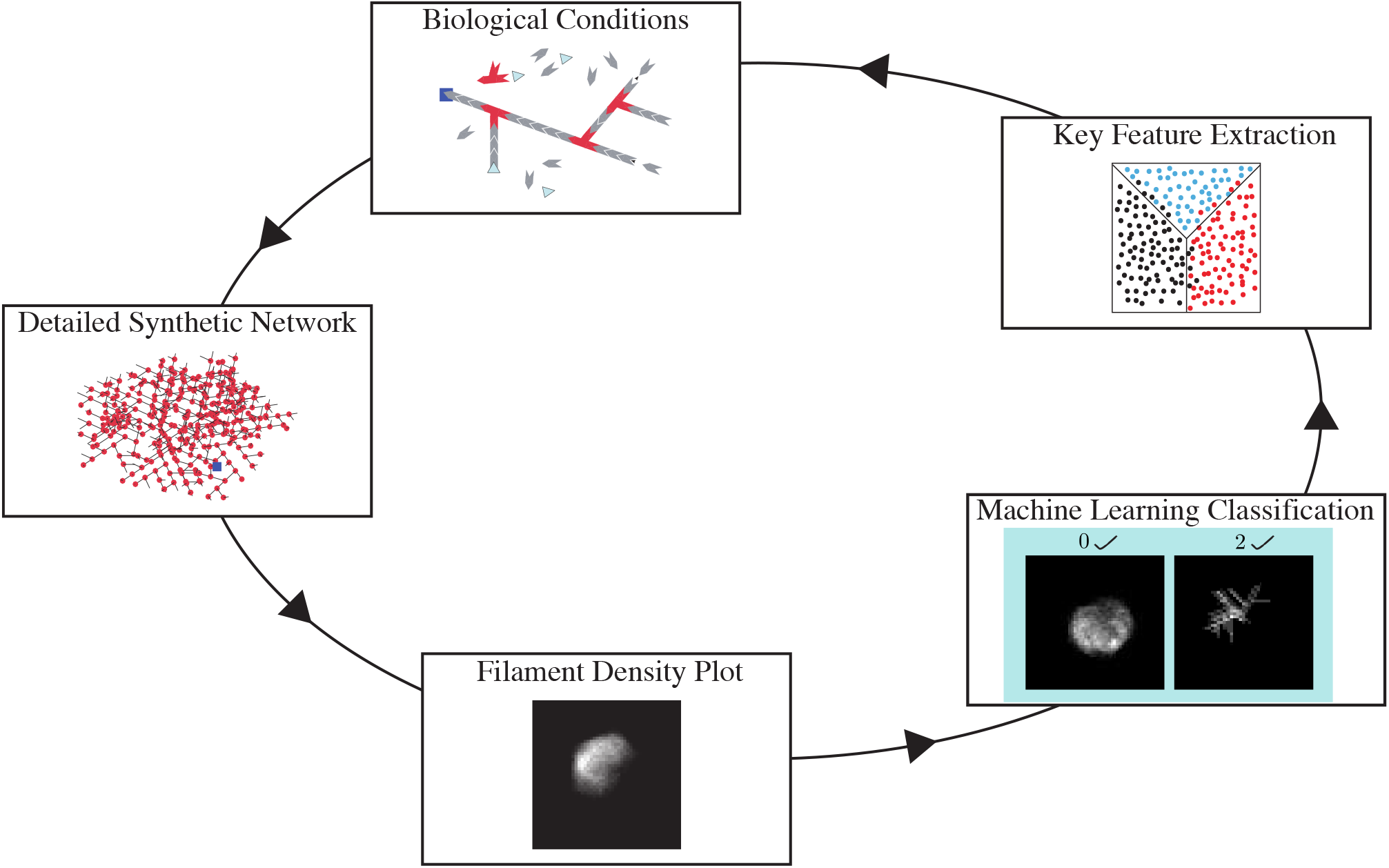
Schematic of the integration of mathematical, computational, and biological techniques. Iteration of these techniques yields insights into mechanisms underlying branched actin network growth.

## Acknowledgments

M.W.R. was supported, in part, by the National Science Foundation (NSF) under Grant No. DMS-1818833 and Grant No. DMS-2146191 (CAREER). K.G. was supported in part by the Natural Sciences and Engineering Research Council of Canada (NSERC) Discovery Grant No. RGPIN-2022-03924. R.L.P. acknowledges support from the Schmidt Science Fellows program, in partnership with the Rhodes Trust. C.C. was supported in part by the NSF under Grant No. DMS-2209494. A.T.D. was supported by the NSF under Grant No. DMS-1554896 (CAREER) and the National Institutes of Health (NIH) under Grant No. R01GM132651.

We would also like to thank the Banff International Research Station (BIRS) for hosting us in May 2022. This BIRS Focused Research Group (FRG) meeting was crucial to the development of this work.

## Appendix

### A.1. Detailed description of stochastic simulation of actin networks

In our previous work, we built a tractable agent-based stochastic model to capture the local microstructure of branching actin networks under various intracellular conditions (7). The model includes the actin dynamics of polymerization, depolymerization, and branching of filaments initiated from a single nucleation site. The actin filaments are represented as rigid rods with a spatially fixed base and a barbed tip capable of growing or shrinking. Changes in both actin filament length and overall network structure are due to the addition or removal of actin monomers and branching via the Arp2/3 complex. In the initial model, we assume that there is an unlimited pool of free monomers and Arp2/3 complexes available to enable filament growth and branching, respectively. For simplicity, other dynamics of actin networks such as capping, sliding, bundling, etc. are not considered in the original model.

Briefly, each simulation starts with an actin filament of length zero located at the “nucleation site” placed at the origin. The filament is assigned an angle of growth which is drawn from a static uniform distribution; the angle of growth prescribes the direction of growth for that particular filament. At each time step, there are four possibilities for filament dynamics: (1) the filament grows with probability 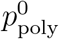; (2) the filament shrinks with probability 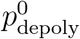, provided that the filament has a nonzero length at the start of the time step; (3) the filament remains the same length; or (4) the filament branches with probability 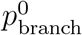 to create a new filament, provided that the original filament is at least some critical length *L*_branch_ (measured from the closest branch point). Filament growth and shrinkage occur in discrete increments corresponding to the length of an actin monomer, Δ*x* 0.0027 *μ*m. Based on the known biological interaction of the Arp2/3 complex with actin, the newly branched filament is assigned to grow in a direction that is±70° from the preexisting filament tip.

To determine which of the four outcomes happens at a time step, two independent uniformly distributed random numbers are generated for each filament in the simulation. If the first random number is less than 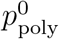, then polymerization occurs, and if it is greater than 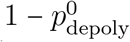, then depolymerization occurs. If the first random number simultaneously satisfies both inequalities, then the filament length is unchanged since both polymerization and depolymerization occur in the same time step. Likewise, filament length is unchanged if neither inequality is satisfied, since this means neither polymerization nor depolymerization occur in the given time step. If the second random number is less than 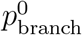, and the filament is at least length *L*_branch_, then the filament branches to form a new filament, capable of autonomous growth and branching. This step-wise process is repeated until the final simulation time is reached. Ultimately, this stochastic model captured molecular-level effects within the network and sensitivity analysis revealed that the biological parameters responsible for filament growth and branching kinetics impacted resulting network dynamics and morphology in a complementary manner. Simulations of branching actin networks were implemented in a custom Matlab code, with additional implementation details available in (7).

In the current study, to capture more biologically relevant branching actin networks, we include capping proteins, limited actin monomers, and limited Arp2/3 branching complexes as follows (schematic in Figure 2A; the full algorithm flow chart is shown in Appendix Figure A1). The capping protein regulates actin polymerization by binding to the barbed end of an actin filament, which blocks the addition and loss of monomers from that. In our simulation, capping dynamics are modeled as follows. At each time step, a third uniform random number is generated for each uncapped filament. If the capping probability (*p*_cap_ = rate of capping × time step) is greater than the random number, then the filament can no longer polymerize or depolymerize. Capping is irreversible; once a filament is capped, it is capped for the remainder of the simulation. In simulations that include limited resources, we start with a fixed number of actin monomers (*M*_0_ = 10, 000) and/or a fixed number of Arp2/3 complexes (*A*_0_ = 24). A polymerization (or branching) event can only occur if there are available actin monomers (Arp2/3 complexes), and the probability of polymerization (branching) depends on how many actin monomers (Arp2/3 complexes) remain. In our initial model with unlimited resources, the polymerization rate was fixed: 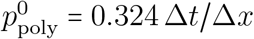, where Δ*t* is the constant time step, Δ*x* is the constant space step. Now, with limited resources, we assume that the polymerization rate depends on the number of actin monomers: 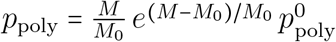, where *M* is the number of available actin monomers at the given time step. Similarly, the branching probability is modified by the number of available Arp2/3 complexes: 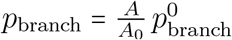, where 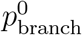 is the branching probability from the unlimited resources case (chosen from a cumulative distribution function of the standard normal distribution with mean 2 and standard deviation 1), and *A* is the number of available Arp2/3 complexes at the given time step. Values for parameters are listed in Appendix Table A1, and explained in detail in the next section.

#### A.2. Stochastic simulation parameters

Wherever possible, model parameters are based on experimental values. The probability of polymerization is calculated from Pollard and Borisy (2003), which states that actin monomers elongate the barbed ends of actin filaments at a velocity of 0.3 *μ*m/s (25). Using the formula

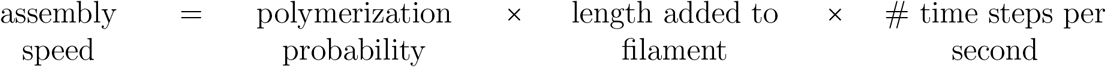

we calculate that

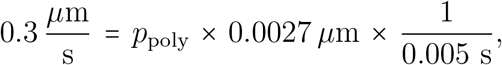

which implies that *p*_poly_ = 0.56 ≈ 0.6. Similarly, Pollard and Borisy (2003) give a range of dissociation rates of actin monomers from 1.4 − 8 s^−1^ (25). We choose an intermediate value of 5.0 s^−1^ and then calculate

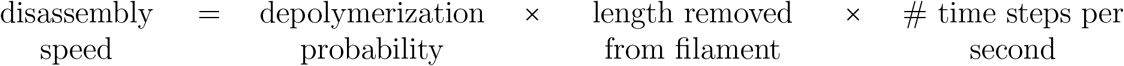

where disassembly speed is obtained via

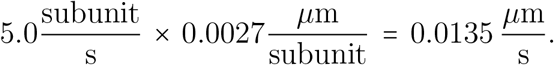

Therefore,

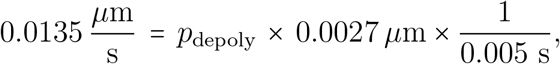

**Figure A1:**
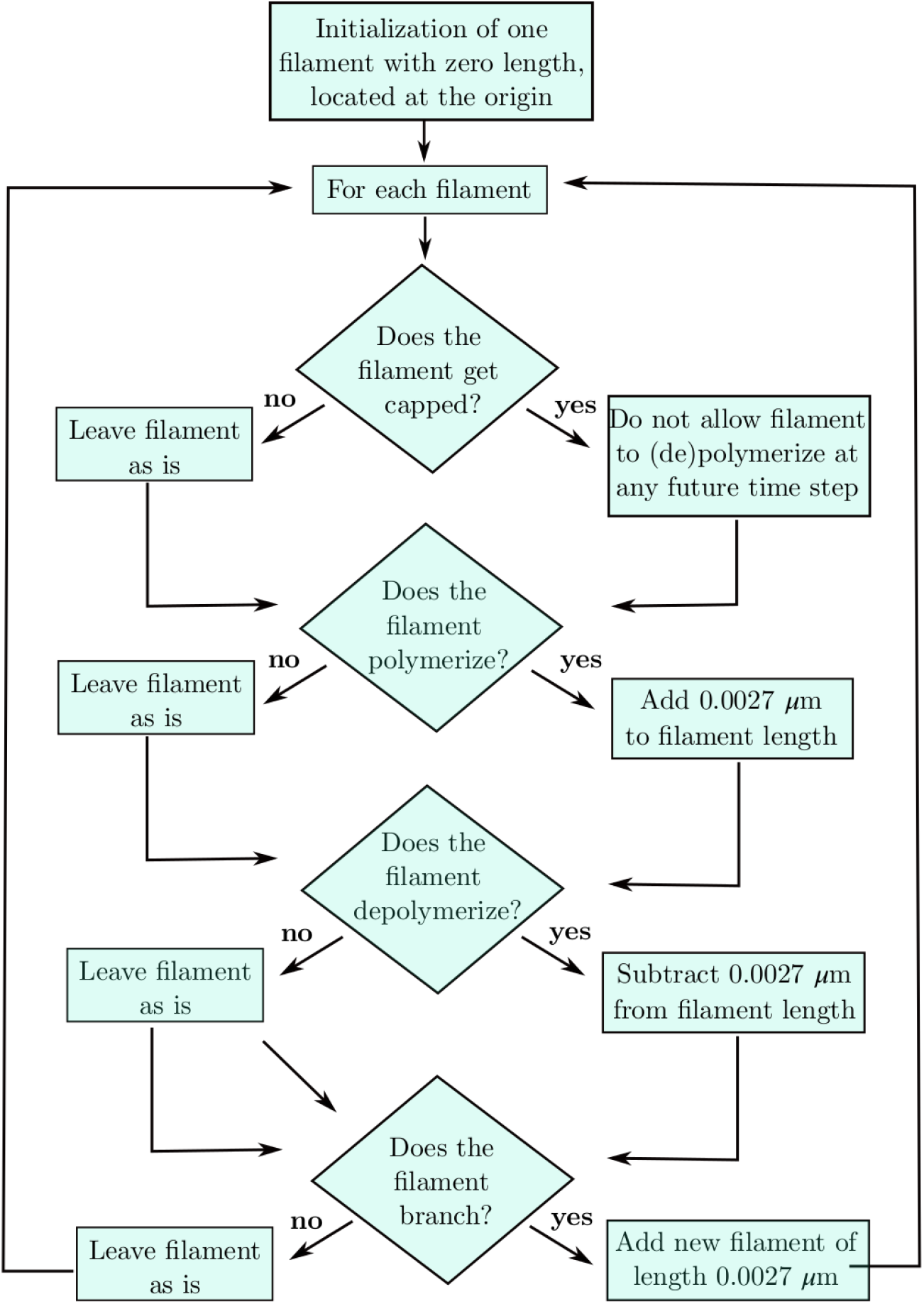
Microscale model flow diagram.

which yields *p*_depol_ = 0.025. Literature measurements of actin filament length per branch vary from 0.02 to 5 *μ*m (26; 6; 27; 28; 29; 30), so we choose *L*_branch_ = 0.2 *μ*m, which is similar to the values from (29; 30) and is an intermediate value between the orders-of-magnitude-different literature values. The probability of capping is calculated using the formula

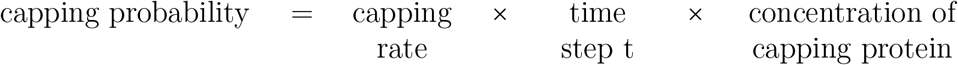

which gives

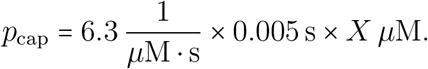

The capping rate of 6.3 *μ*M^−1^s^−1^ is from (31), and *X*, the concentration of capping protein, can take on values up to 0.168 *μ*M (32), which corresponds to *p*_cap_ = 0.0053. Hence, we look at a range of capping probabilities from 0 to 0.00375; capping probabilities greater than this do not result in appreciable actin network growth.

**Table A1:**
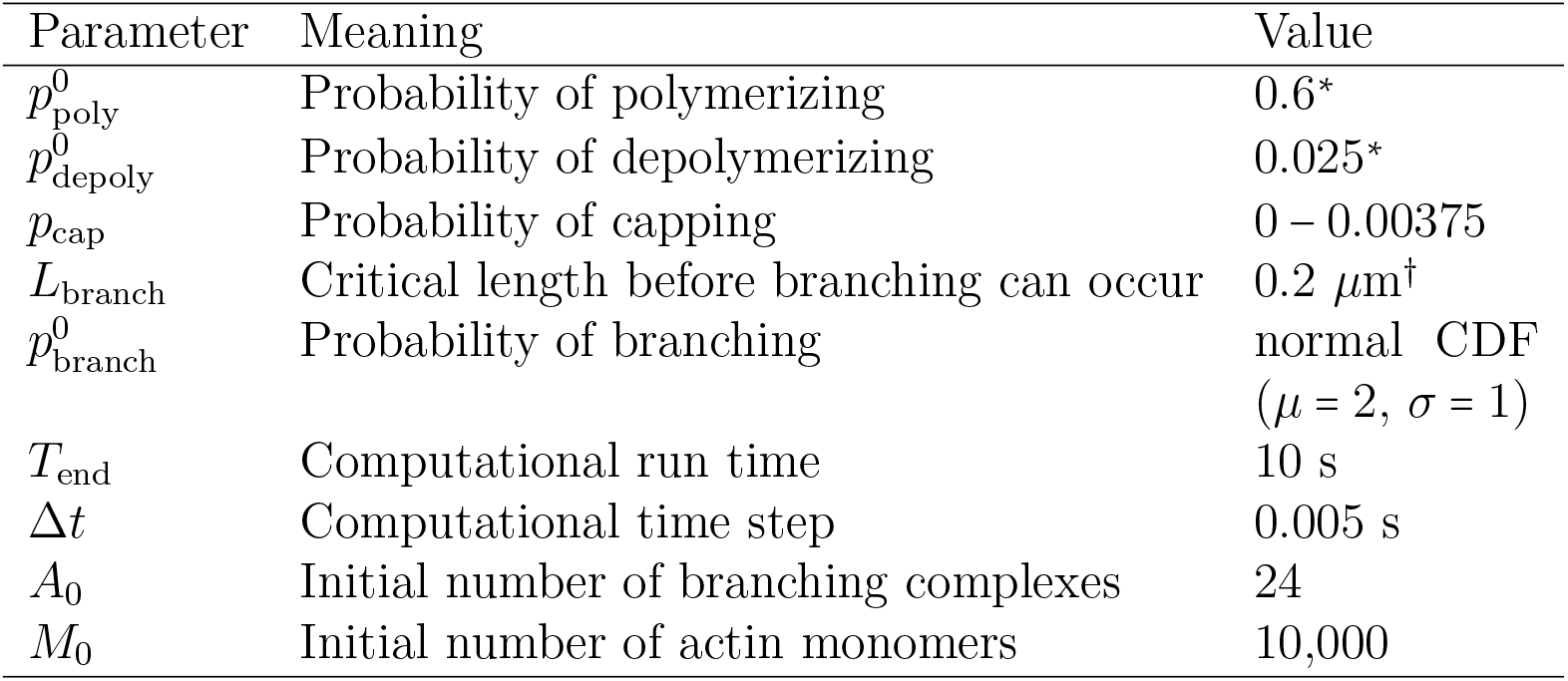
Microscale model parameter values. Values flagged with one star (^*^) were calculated from (25). For the value flagged with a dagger (^†^): literature measurements of actin filament length per branch vary from 0.02–5 *μ*m (26; 6; 27; 28; 29; 30). We choose *L*_branch_ = 0.2 *μ*m as an intermediate estimate between these orders-of-magnitude-different values from the literature, similar to the values from (29; 30). 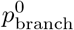 is chosen from a Gaussian distribution with mean *μ* = 2 and standard deviation *σ* = 1. A range of *p*_cap_ values from 0 to 0.00375 are investigated.

#### A.3. Scaling

For visualization purposes, a grayscale image can be created for each actin network based on its filament density. More precisely, the location with the highest density is assigned a 1 (brighest), and the location with the lowest density is assigned a 0 (darkest). As the density varies considerably among networks generated under the eight different growth conditions, this re-scaling is performed network by network instead of across all networks to ensure that networks of lower densities are still visible. Figure A2 shows the re-scaled density plots of two networks generated under Condition 0 (no capping with unlimited Arp2/3 complexes and actin monomers) and Condition 7 (capping with limited Arp2/3 complexes and actin monomers). These two conditions produce the densest and sparsest networks. By comparing the two plots, we see that the same level of brightness can indicate vastly different densities. For all computational purposes, such as the training and testing of classifiers, we use the raw, un-scaled actin density.

**Figure A2:**
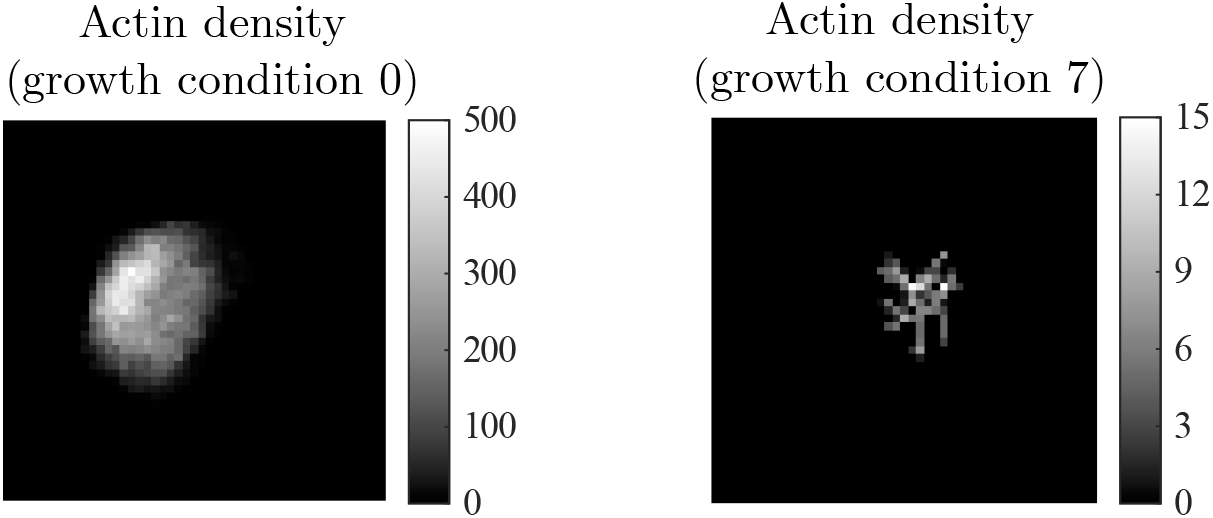
Comparison of the grayscale images of the density of two actin networks, generated under growth condition 0 and growth condition 7 respectively, when the capping probability is 0.001.

#### A.4. Sensitivity to rotation

To determine if our classification method is sensitive to rotation of images, we rotate all density plots in the training and test data sets for capping probability 0.001 clockwise by uniformly random degrees between 0 and 360. Sample rotated density plots (corresponding to the density plots from main text Figure 4) are shown in Figure A3. We retrain the fine-grain CNN classifier based on this rotated training set, and then classify the rotated test set. The accuracy of the fine-grained CNN classifier on the rotated images is 89%, identical to the accuracy of the method on the original images. Hence, we conclude that our method is not sensitive to rotation.

**Figure A3:**
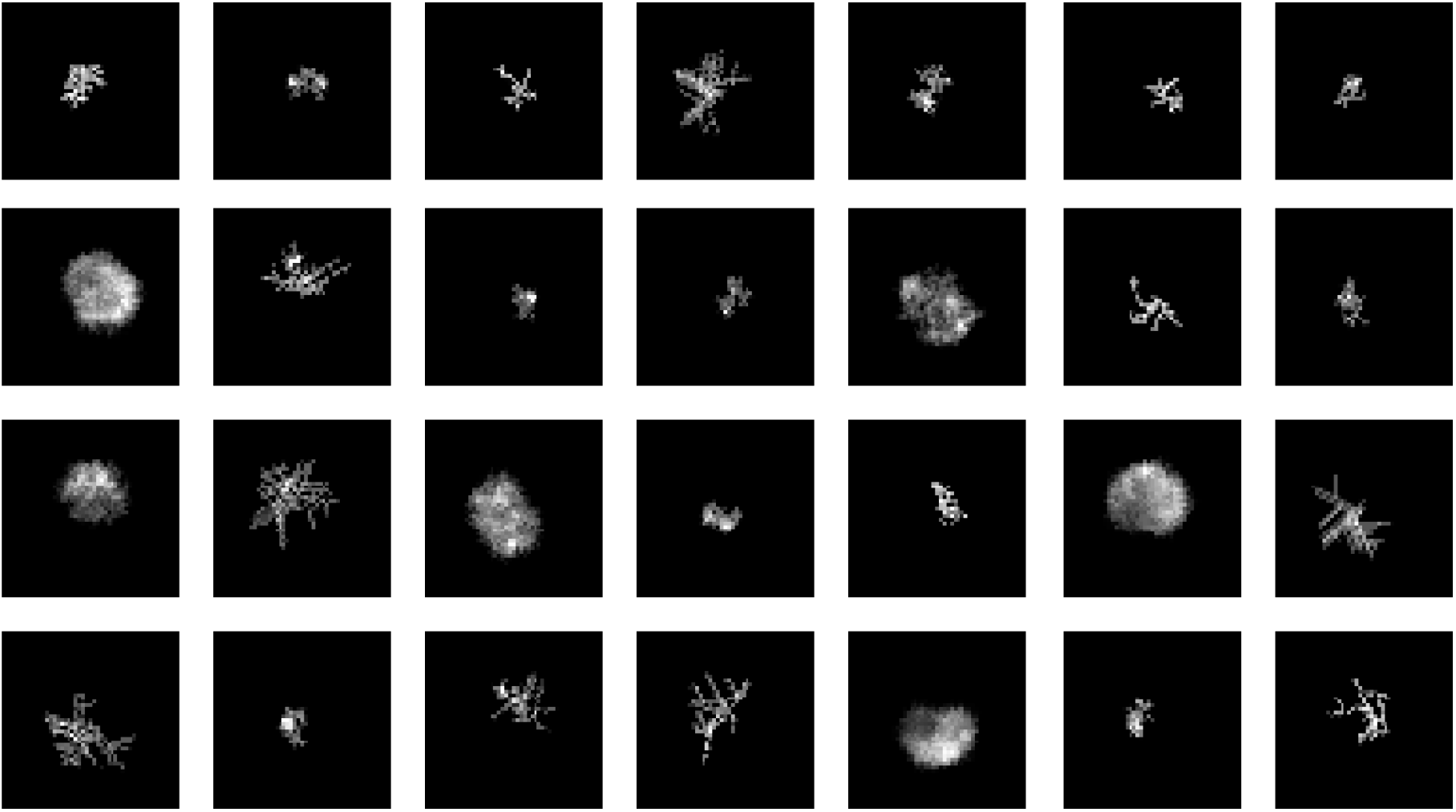
Samples of rotated density plots from the training and test sets.

#### A.5. Sensitivity to salt and pepper noises

To determine if our method is sensitive to noisy data, we “pollute” with salt and pepper noise the density plots in the training and test sets for capping probability 0.001. In our first experiment, we randomly select 1% of pixels and randomly change each one to white or black. We then retrain the fine-grain CNN classifier on the noisy training set, and then classify the noisy test set. The accuracy of our method with 1% of pixels randomly changed is 85% (compared to 89% with the original, non-noisy images). We repeat this process with 5% and 10% of pixels randomly set to white or black and obtain 85% and 82% accuracy, respectively. Sample density plots (corresponding to the density plots from main text Figure 4) with 10% of salt and pepper noise added are shown in Figure A4. We conclude that our method is not very sensitive to noise.

**Figure A4:**
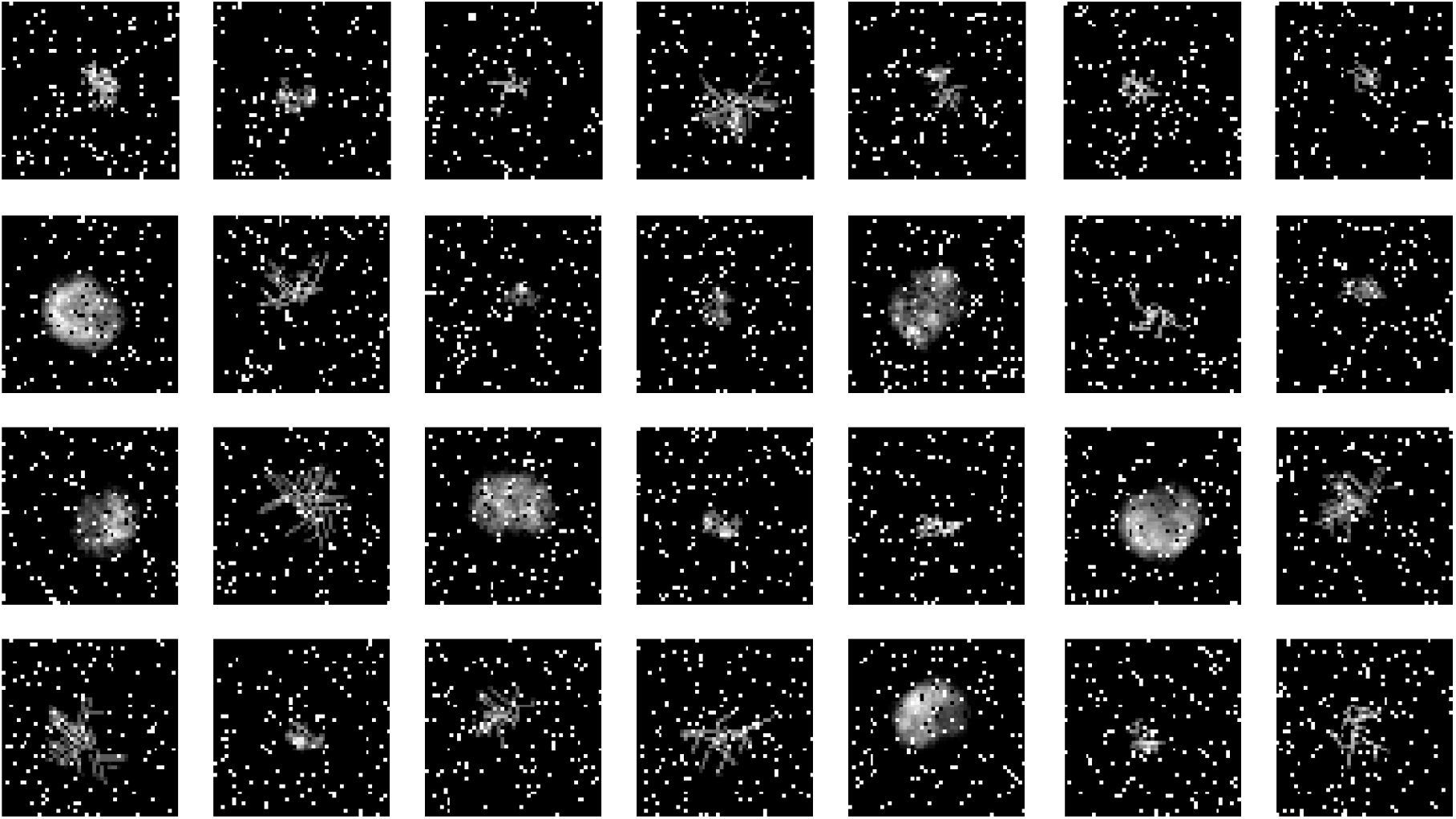
Samples of density plots from the training and test sets with 10% of salt and pepper noise added.

#### A.6. Sensitivity to time of data collection

To determine if our method is sensitive to the time of data collection, for capping probability 0.001 we generate test sets at final simulation times *T* = 5, 7, 8, 9, 9.5, 9.8, 10.5, 11, and 12 seconds and classify them using the fine-grain CNN classifier obtained based on training set data collected at time *T* = 10 s. Snapshots of a single actin network at the different times are shown in Figure A5. Our method is somewhat sensitive to time of data collection (Figure A6). The accuracy of the method is best (89%) when both the training and test sets are collected at time *T* = 10. The method is reasonably accurate for test sets collected at *T* = 9.5, 9.8, 10.5, 11, and 12 seconds (87%, 89%, 89%, 86%, 79%, respectively), but not very accurate (accuracy < 75%) for test sets collected at *T* values less than or equal to 9 s. Despite the sensitivity of the method at early time points, we conclude that the classifier is robust enough for some human error, since it gives accurate classifications for test sets collected between 9.5-12 seconds.

**Figure A5:**
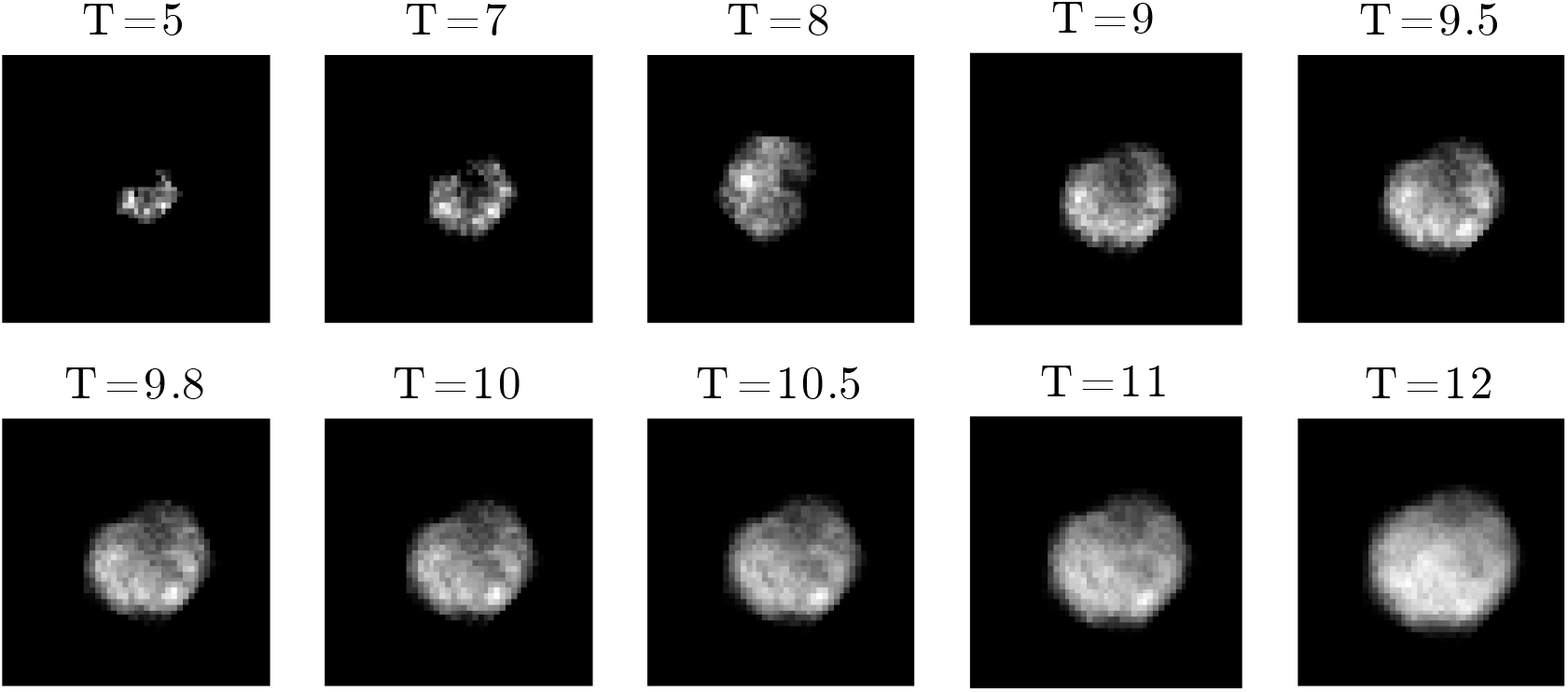
Snapshots of an actin network from the test set at various times.

**Figure A6:**
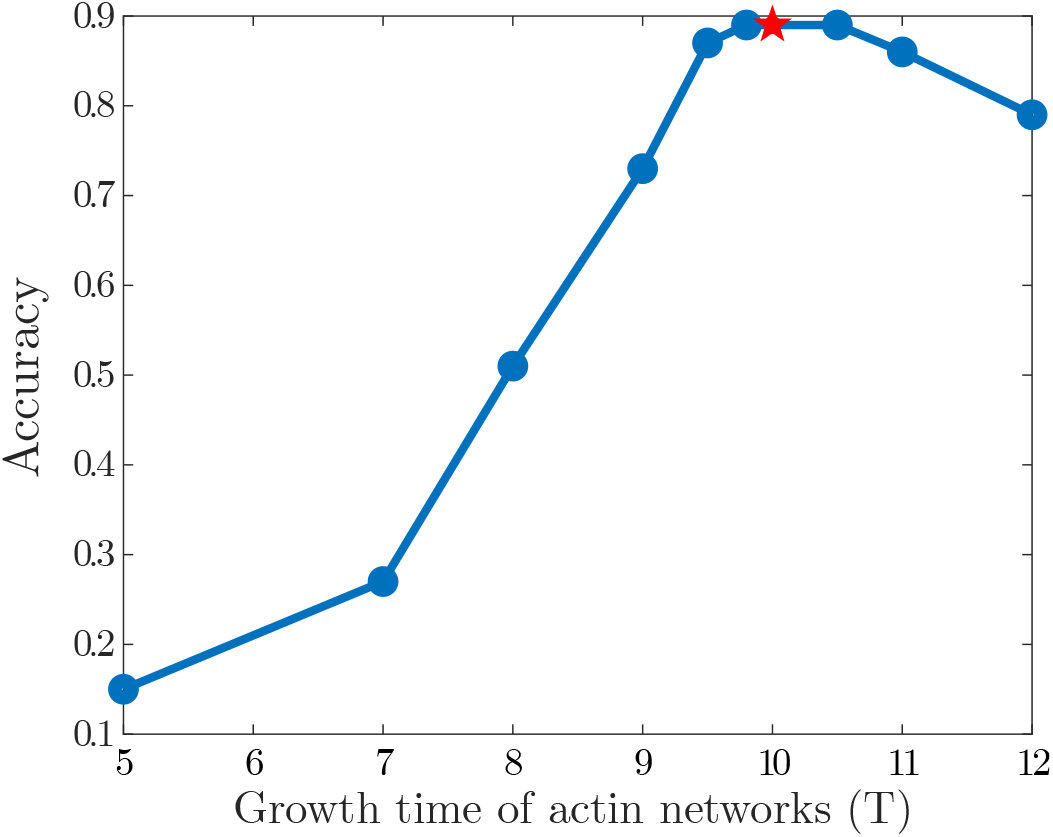
Accuracy of the fine-grained CNN classifier for time *T* = 10 on test sets obtained at various time points.

## Footnotes

1 First author

2 https://github.com/bbannish/actin

3 https://www.mathworks.com/help/stats/classificationlearner-app.html

4 https://www.mathworks.com/help/deeplearning/index.html

5 https://www.mathworks.com/help/stats/index.html

6 https://www.mathworks.com/help/stats/index.html

